# Genetic drivers of protein changes over time: Findings, considerations, and approaches in TOPMed cohorts and UK Biobank

**DOI:** 10.64898/2026.08.03.742409

**Authors:** Madeline G. Gillman, Han Chen, Annie Green Howard, Michael Mi, Zsu-Zsu Chen, Clary B. Clish, Daniel E. Cruz, Peter Durda, Craig Johnson, Ani W. Manichaikul, Suna Onengut, Prashant Rao, Usman A. Tahir, Kent D. Taylor, Russell P. Tracy, Alexis C. Wood, Robert E. Gerszten, Lifang Hou, Ravi Shah, Jerome I. Rotter, Stephen S. Rich, Laura M. Raffield

**Author notes:** Correspondence: Laura Raffield, Department of Genetics, 120 Mason Farm Road, 5042 Genetic Medicine Building, #7264, Chapel Hill, NC, 27599, USA.

## Abstract

Age is a major risk factor for many diseases, but the biological processes driving aging are heterogeneous across individuals. Efforts to untangle differences between chronological and biological age have focused on identifying age-associated markers, such as ’omics clocks. Many ’omics features, including proteins, are strongly associated with age, and genetics contribute to variance in these measures. However, few studies have identified genetic drivers of interindividual variability in ’omics changes over time.

Using longitudinal proteomics data (Olink 3k) from the Multi-Ethnic Study of Atherosclerosis (MESA), we calculated a protein slope for each individual (n=2,007) and protein (n=2,737) across 3 visits spanning 14–18 years, then conducted a genome-wide analysis for each slope, both with and without adjusting for baseline protein level. Subsets in UK Biobank (UKB; n=948) and CARDIA (n=1,328) with longitudinal proteomics data were used for replication. We considered additional methods for modeling of protein change and variability, including linear mixed models, SNP-by-age interactions, and variance quantitative trait loci.

Without baseline adjustment, only 19 proteins (20 credible sets) had a slope pQTL in MESA, with poor replication in UKB and CARDIA. With baseline adjustment, 607 proteins (698 credivle sets) had a slope pQTL and over 70% replicated in CARDIA and/or UKB; such baseline adjusted models may, however, be subject to collider bias. Longitudinal and cross-sectional interaction models identified fewer than 14 pQTLs, suggesting they were generally underpowered; but 73% of proteins with a variance pQTL also had a slope pQTL.

By examining effect direction concordance, replication rate, directed acyclic graphs, and signal overlap with other models we demonstrate that many baseline-adjusted slope pQTLs may be arising due to model misspecification or regression to the mean. Overall, our results highlight considerations for modeling strategies of change phenotypes and build on understanding of potential genetic mechanisms influencing interindividual proteome changes over time.

## Introduction

Physiological changes as humans age are profound and age itself is a major risk factor for many diseases(1). Rate of aging varies, and there is an increasing interest in identifying the mechanisms driving differences in biological and chronological age. As the body’s main molecular workhorse, protein measures, especially from blood (which contains proteins from almost all body tissues), are an apt lens through which to view how changes associated with age occur, including variation across individuals in these age-related changes. Attempts to untangle age-related changes have primarily focused on identifying age-associated signatures or using machine learning methods to create aging “clocks” using various ‘omics measures.(2,3) The genetic underpinnings driving differences in rates of change of individual proteins has yet to be explored.

There has been an increasing interest in using proteomics to identify mechanisms and biomarkers associated with biological age. Proteins associated with age in the form of aging clocks have been identified,(3,4) but not as much work has described within-individual changes in protein levels over 10+ years. Over shorter periods of time (e.g. 2 years), proteomics measures are relatively consistent(5–7) but measures do change over longer periods: a longitudinal study measuring 84 proteins in ∼1,000 participants found that over 80% of proteins significantly change within individuals over a period of 10 years, with most proteins measured on this highly targeted panel showing an increase in abundance(8). Thousands of genetic variants have been associated with protein abundance measures in cross-sectional pQTL studies. Proteins can also vary differently across genotypes, suggesting some level of environmental interaction. Studies in UK Biobank have identified hundreds of variance pQTLs (which can be used as a proxy for gene-environment interactions, indicating that the genetic regulation of protein levels is modified by exposures).(9,10) While intra- and inter-individual changes in protein expression across age have been demonstrated, many of the genetic mechanisms governing these changes have not been identified.

Prior work has demonstrated that trait changes can have genetic associations (e.g., with BMI(11,12), systolic blood pressure(13), and eGFR(14)), and that using the trajectory, change, or slope measure can improve risk prediction. For example, in a cohort of Black adults (the Jackson Heart Study), the baseline measure of C-reactive protein (an inflammatory marker protein that is a predictor of cardiovascular diseases(15)) was not associated with incident HF, but changes in CRP from high-to-low and low-to-high were associated with incident HF(16). Trajectories of other common disease risk phenotypes such as eGFR(14), systolic blood pressure(13,17) BMI(11,12), and cholesterol(17) have been linked to genetic variants, suggesting that the rate of change in these phenotypes has in part a genetic basis. The rate of change of a protein has also been associated with health outcomes; for example increases in NT-proBNP have been found to be predictive of cardiovascular events(18), and changes in the proteome linked to severity of COVID-19(19).

Innovations in high-throughput proteomics have enabled the study of a large fraction of the circulating human proteome. While studies using cross-sectional measures have revealed genetic insights, but the use of these assays in a longitudinal context is newer. The semi-quantitative nature of these assays raises questions of how data transformation impacts longitudinal modeling, and the extraordinary number of features combined with computational burden of common longitudinal analysis techniques necessitates careful modeling choices. Traits with known rates of change, such as eGFR (which declines at an approximate rate of -1 mL/min/1.73 m^2^ per year starting at around age 40), have enabled deeper analysis to select methods that best model the ground truth rate of change(14). Cases where only one trait is of interest, and more observations per participant are available (e.g. with BMI), enable more complex modeling strategies—e.g. genome wide longitudinal GWAS.(12,20)

In this work, we have three aims: First, we compare strategies for determining a protein rate of change using Olink 3k data from three visits in the Multi-Ethnic Study for Atherosclerosis (MESA). Second, we then use our selected method to conduct a genome-wide slope pQTL analysis (**Figure 1**) and characterize slope loci using longitudinal Olink data from the MESA, Coronary Artery Risk Development in Young Adults (CARDIA), and UK Biobank (UKB) studies, including exploring the impact of adjusting for the baseline protein measure. Third, we explore slope pQTL overlap with more complex modeling methods including cross-sectional and longitudinal SNPxAge interaction models and two-stage random slope random intercept models. We additionally explore overlap with cross-sectional variance pQTLs(10) to improve understanding of the potential overlap between genetic regulation of longitudinal protein changes and cross-sectional protein variability. Studying protein slopes can provide insight into the molecular mechanisms governing biological traits and could identify loci with greater clinical potential in understanding of future incident disease than those associated with single timepoint measures alone, thus helping to refine prediction models and personalized medicine approaches.

**Figure 1.**
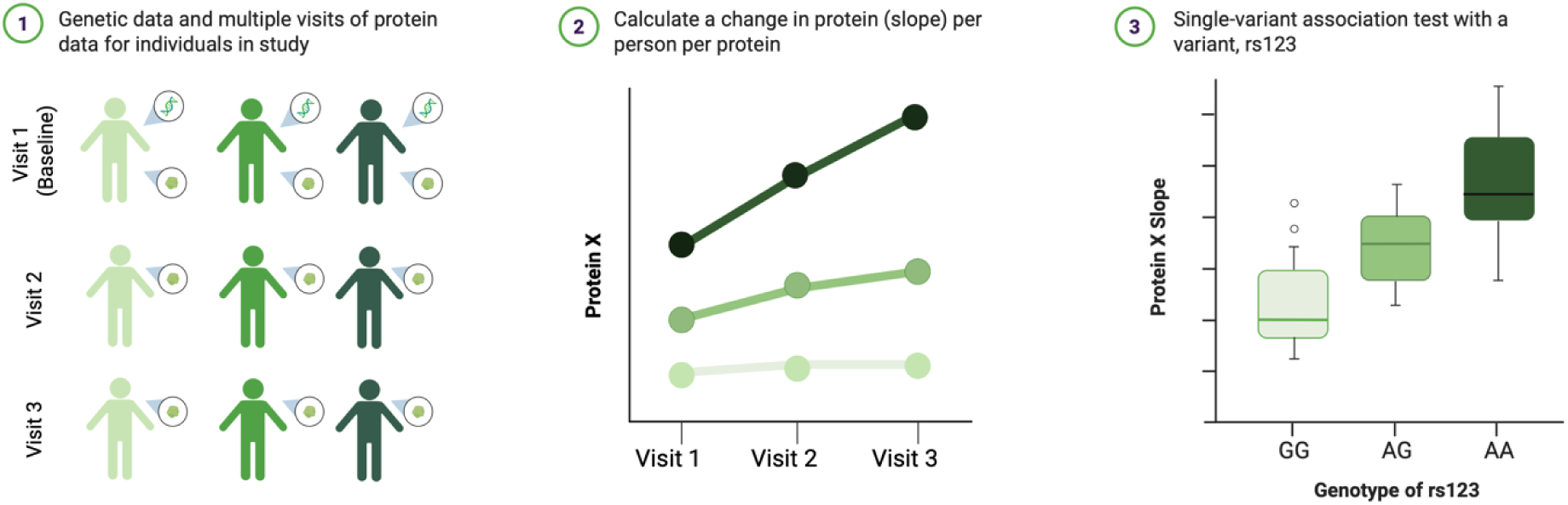
Cartoon example of a slope pQTL. In this example, the A allele in rs123 is associated with a greater slope of Protein X.

## Results

### Demographics

On average, protein slopes in MESA captured change over 12 years of follow-up, 14 years in CARDIA, and 12 years in UK Biobank (Table 1). The average age at baseline for MESA participants was 58 years while CARDIA and UKB were generally younger (40 years in CARDIA and 50 years in UKB).

**Table 1.**
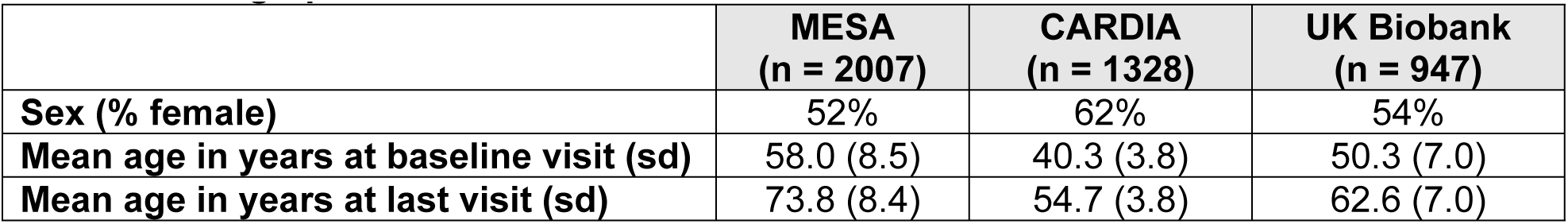
Demographic characteristics of cohorts used.

|  | MESA<br>(n = 2007) | CARDIA<br>(n = 1328) | UK Biobank<br>(n = 947) |
| --- | --- | --- | --- |
| Sex (% female) | 52% | 62% | 54% |
| Mean age in years at baseline visit (sd) | 58.0 (8.5) | 40.3 (3.8) | 50.3 (7.0) |
| Mean age in years at last visit (sd) | 73.8 (8.4) | 54.7 (3.8) | 62.6 (7.0) |

### Proteomics data and transformations

Because each cohort was run on a different version of the Olink platform, protein overlap varied. After dropping proteins with high missingness and high CVs in each cohort, 1,922 were measured in MESA and CARDIA, 1,453 in MESA and UKB, and 1,132 proteins were measured in all three cohorts (as determined by UniProt IDs and matching Olink IDs if applicable).

Pairwise comparisons of protein slopes calculated in MESA from each of the three NPX data transformations revealed that resulting slopes were generally similar regardless of the transformation applied (**Figure 2**). Pairwise correlations between protein means, slopes, and each cross-sectional measure reflected expected relationships (**Supplemental Figure 1, Supplemental Figure 2A).** Slopes calculated using three visits and two visits (essentially a difference model) were highly correlated (Pearson r=0.99, **Supplemental Figure 2**).

**Figure 2.**
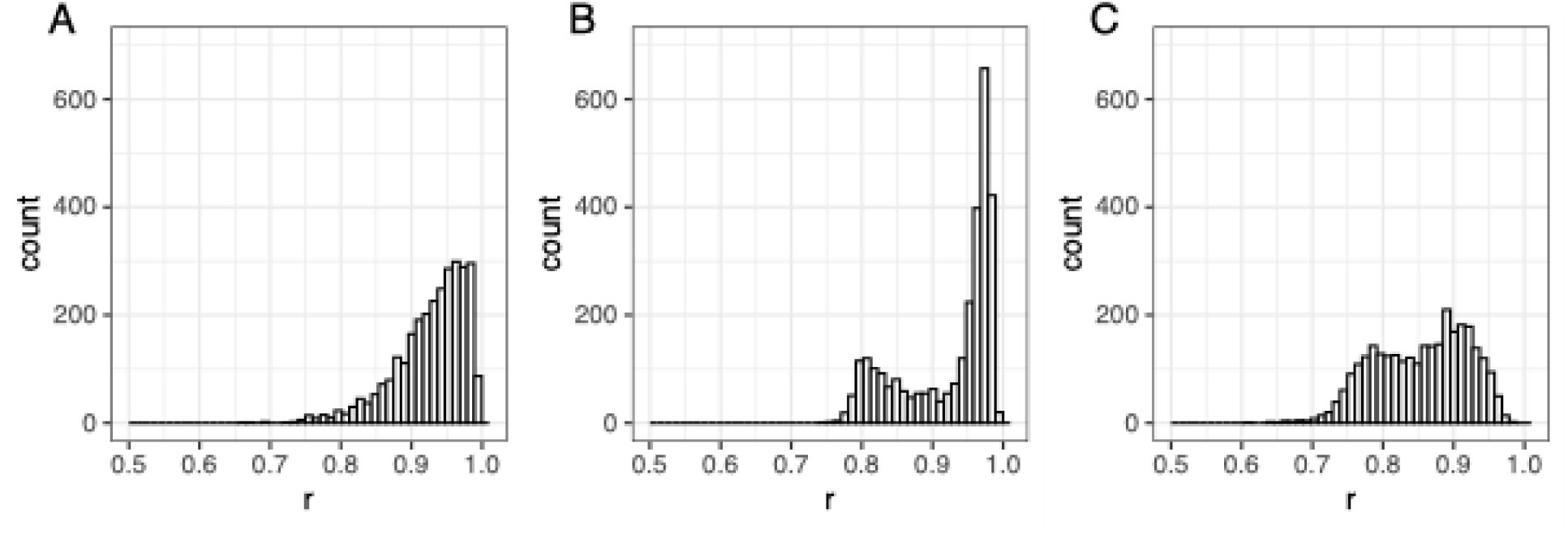
Different data transformations prior to calculating slope result in similar slope values. **(A)** Histogram of Pearson’s correlation coefficients comparing a linear slope calculated using the NPX values (which have been either intensity or plate control normalized depending on protein) versus the INT values (which have been scaled and inverse normal transformed within visit prior to the slope calculation) (median correlation = 0.94). **(B)** Histogram of Pearson’s correlation coefficients comparing a linear slope calculated using the protein residuals (obtained by regressing the protein values on age, sex, site, and plate within visit) versus the INT values (median correlation = 0.95). **(C)** Histogram of Pearson’s correlation coefficients comparing a linear slope calculated using the protein residuals (obtained by regressing the protein values on age, sex, site, and plate within visit) versus a slope calculated using the NPX values (median correlation = 0.86).

### Slope pQTL discovery in MESA

Using the baseline unadjusted model only 19 proteins had a slope pQTL, resulting in 20 credible sets (**Figure 3C-E**). Only one protein had more than one credible set identified (CPXM1) and only one locus (for SNAP25) contained a confidently fine-mapped likely causal variant (top variant PIP >= 0.95). Effect directions were generally discordant compared to the cross-sectional effects, e.g. for the SNAP25 protein in **Figure 4**.

**Figure 3.**
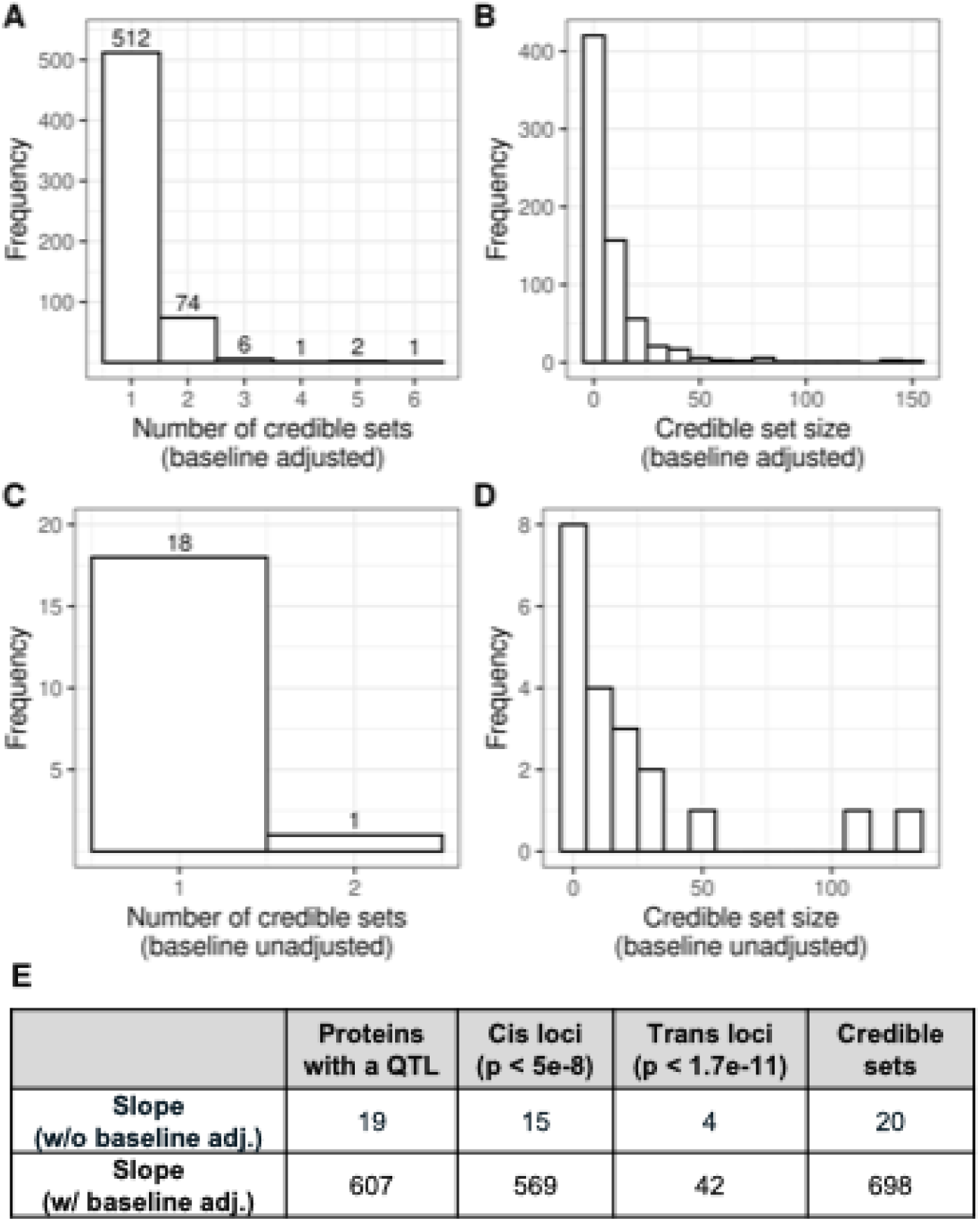
Slope pQTL credible set summary. Credible sets were identified using the summary statistics from the baseline-adjusted analyses with SuSiE using individual-level LD. SuSiE failed to identify a credible set for three proteins in the baseline adjusted analysis. Distribution of number of credible sets identified per protein in MESA in baseline adjusted analysis (**A**) and baseline unadjusted analysis (**C**). Distribution of number of variants per credible set in baseline adjusted analysis (**B**, note that one outlier credible set with 1,907 variants is excluded) and baseline unadjusted analysis (**D**). (**E)** Number of loci identified in MESA in baseline adjusted and baseline unadjusted analyses. Loci were identified using distance-based clumping with a 1Mb window. Credible sets were identified with SuSIE using individual-level LD.

**Figure 4.**
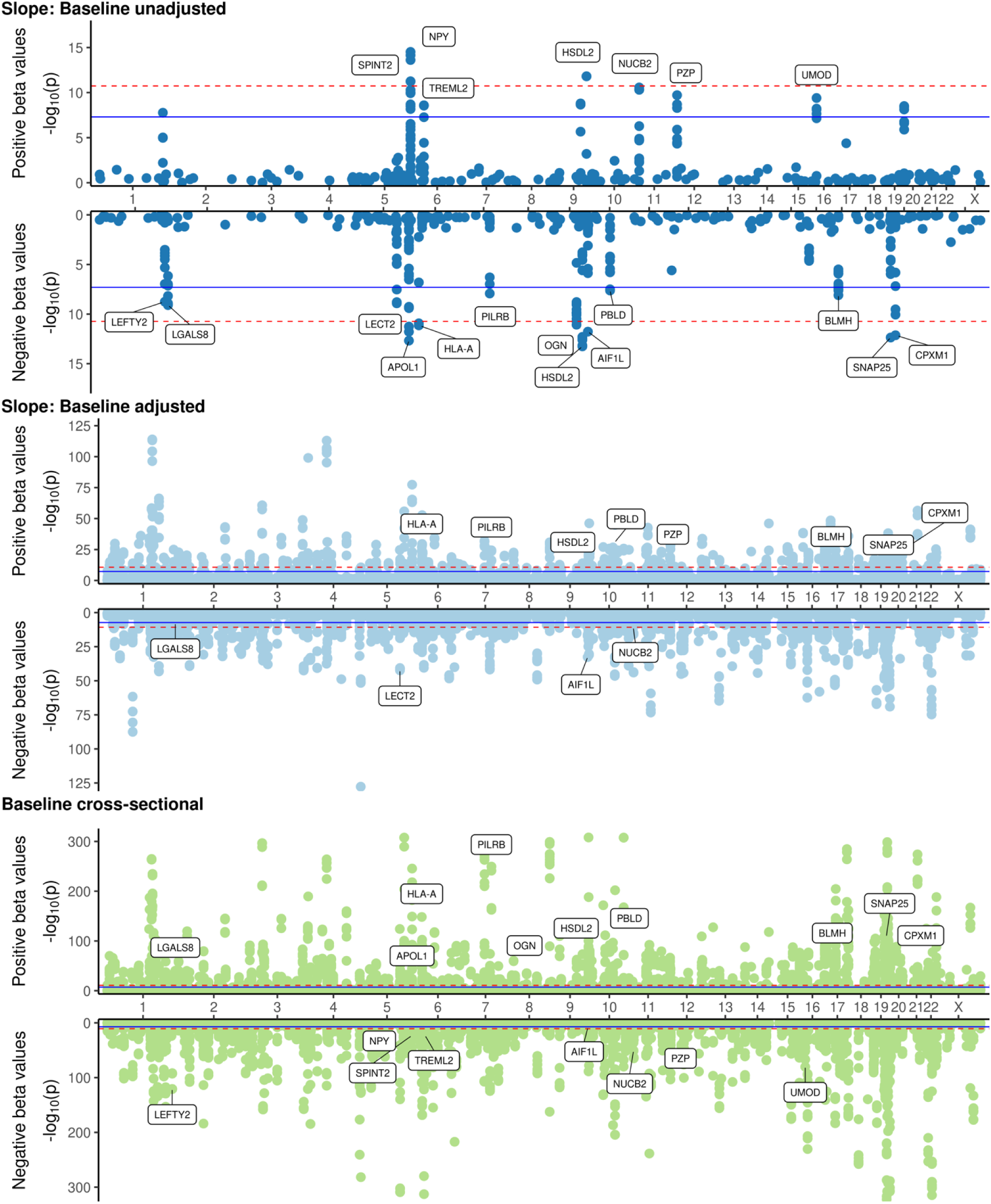
Miami plots of significant loci in slope baseline unadjusted, slope baseline adjusted, and cross-sectional analyses. Only significant loci in the baseline unadjusted model are labeled in each plot. For the baseline cross-sectional plot, data plotted only represents proteins with a slope pQTL (either baseline unadjusted or baseline adjusted).

Using the baseline adjusted model, 607 proteins from 611 loci (based on distance-based clumping) had a slope pQTL identified in MESA (**Figure 3E**). SuSiE identified 698 credible sets from 596 proteins; visual inspection of loci where fine-mapping failed to converge showed that they were often in complex LD regions or had a p-value near the significance threshold. The majority of proteins had only one credible set identified with a mean credible set size of 12.8 (range: 1, 1,907) (**Figure 4A, B**). For 191 of the credible sets we were able to determine the likely causal variant (top variant PIP >= 0.95). The majority (over 90%) of the slope pQTL credible sets contained at least one variant that was directionally concordant and significant in the cross-sectional analyses (P_Visit1_ < 5e-8) (**Figure 5, Supplemental Table 2**). Betas in the baseline adjusted analysis were consistently larger and standard errors were consistently smaller than in the baseline unadjusted analysis (**Supplemental Table 2** and **Supplemental Table 3**).

**Figure 5.**
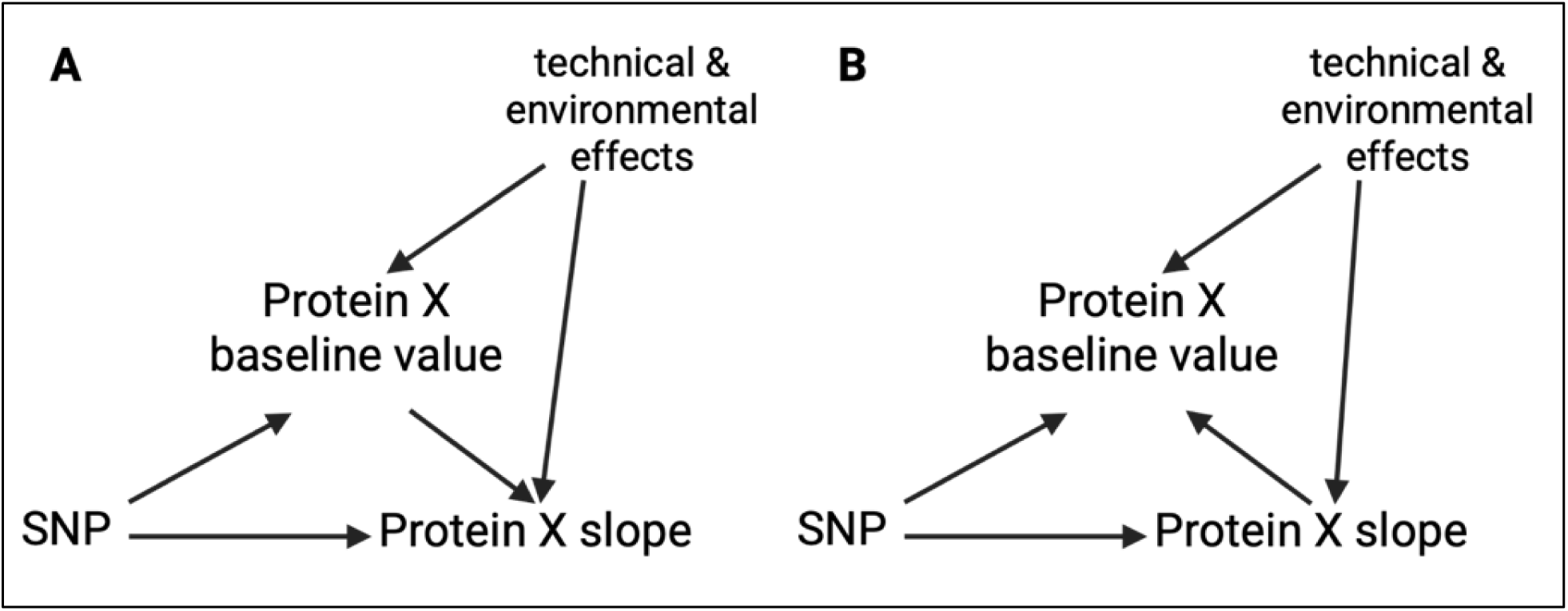
Directed Acyclic Graph (DAG) illustrating potential relationship between SNP, baseline protein value, and protein slope. The baseline measure can either be a mediator as shown in **A,** or it can act as a collider as shown in **B**. With the scenario presented in A, the baseline measure should generally not be adjusted for unless the interest is in the direct effect only. With the scenario presented in B, conditioning on the baseline value would introduce an open backdoor path with the technical and environmental effects.

Baseline adjusted and unadjusted credible sets with lookups from the cross-sectional analysis are presented in **Supplemental Table 2** and **Supplemental Table 3.**

### Comparing slope loci identified in baseline adjusted vs. baseline unadjusted analyses

When comparing the genetic architecture of slope loci in baseline adjusted and baseline unadjusted analyses there is a stark contrast in the number of loci identified and the directionality of signals. The majority of baseline unadjusted loci are directionally discordant with the corresponding baseline adjusted SNP effect and the corresponding cross-sectional SNP effect (e.g., the SNAP25 locus on chromosome 19 in **Figure 4**). On the other hand, the majority of baseline adjusted loci are directionally concordant with the corresponding cross-sectional SNP effect estimate. Overall, credible sets were unique to the model used: only two credible sets identified in the baseline unadjusted analysis comprised SNPs that were also in credible set in the baseline adjusted analysis (**Supplemental Figure 5 and Supplemental Figure 6**). When simply using lookups, the majority of credible sets identified in the baseline unadjusted analysis were unique compared to the baseline adjusted model (i.e. the baseline unadjusted credible sets did not contain any SNPs with P_baseline adjusted_ < 5e-8) and over half of those were directionally discordant with the baseline adjusted model.

To further assess these relationships, credible sets from the baseline adjusted and baseline unadjusted analyses in MESA were grouped into four categories based on SNP effect directionality and significance; then replication was assessed for each category. A Bonferroni-adjusted p-value for the number of MESA baseline unadjusted and baseline adjusted credible set SNPs was used to determine replication (p < 5.31E-06 [0.05/9409 SNPs]). Credible sets that were both significant in the baseline adjusted only model and directionally discordant with baseline unadjusted estimates had the highest replication rate: 93.1% replicated in UKB, and 81.9% replicated in CARDIA (**Table 2**). None of the credible sets identified using the baseline unadjusted model replicated in baseline unadjusted analyses in UKB; however three of the four credible sets that were directionally concordant but not significant in baseline adjusted analyses replicated in CARDIA (**Table 2**).

**Table 2.** Number of credible sets from baseline adjusted and unadjusted analysis that are significant and directionally concordant in baseline unadjusted analysis and their respective replication rates. Baseline adjusted analyses had a total of 698 credible sets identified from 607 unique proteins. Baseline unadjusted analyses had a total of 20 credible sets identified from 19 unique proteins. Credible sets were considered replicated if any of the variants in the credible set (as identified in MESA) had P < 5.31E-06 in the indicated cohort using the corresponding model. A total of 1,404 proteins were measured in both MESA and UKB; 1,865 proteins were measured in both MESA and CARDIA. Replication rates consider only proteins measured in both cohorts specified.

| Category | N proteins (MESA) | N slope pQTL credible sets (MESA) | % replicated in UKB | % replicated in CARDIA |
| --- | --- | --- | --- | --- |
| **Baseline unadjusted analysis** |  |  |  |  |
| Significant in baseline unadjusted and baseline adjusted, directionally concordant | 2 | 2 | NA (proteins not measured in UKB) | 0% (0/2 credible sets) |
| Significant in baseline adjusted and baseline unadjusted, directionally discordant | 4 | 4 | 0% (0/2 credible sets) | 25% (1/4 credible sets) |
| Significant in baseline unadjusted only, directionally concordant with baseline adj. | 4 | 4 | 0% (0/3 credible sets) | 75% (3/4 credible sets) |
| Significant in baseline unadjusted only, directionally discordant with baseline adj. | 9 | 10 | 0% (0/8 credible sets) | 0% (0/10 credible sets) |

| **Baseline adjusted analysis** |  |  |  |  |
| --- | --- | --- | --- | --- |
| Significant in baseline adjusted and baseline unadjusted, directionally concordant | 2 | 3 | NA (proteins not measured in UKB) | 33% (1/3 credible sets) |
| Significant in baseline adjusted and baseline unadjusted, directionally discordant | 2 | 2 | NA (proteins not measured in UKB) | 100% (2/2 credible sets) |
| Significant in baseline adjusted only, directionally concordant with baseline unadjusted | 274 | 311 | 58.3% (81/139 credible sets) | 56.7% (166/293 credible sets) |
| Significant in baseline adjusted only, directionally discordant with baseline unadjusted | 348 | 382 | 93.1% (202/217 credible sets) | 81.9% (289/353 credible sets) |

### QTL discovery using other methods

With a baseline cross-sectional SNPxAge interaction model none of the MESA slope leads or the UKB cross-sectional leads tested had a significant SNPxAge interaction (**Table 3**) in MESA. However the longitudinal SNPxAge interaction model did identify significant associations: 31 of the MESA slope pQTL leads (including baseline adjusted and baseline unadjusted) had a significant age interaction effect; of those 17 were baseline unadjusted slope pQTLs. Of the 20,344 protein-SNP pairs from UKB that we were able to test in MESA with a longitudinal AgexSNP interaction model, 36 had significant age interactions (p < 2.46E-06 [0.05/20,344]) 3 associations were directionally concordant with the main effect (which represents the genetic effect on mean proteins values for an individual with age = 50) and 33 were directionally discordant with the main effect. Summary statistics for the tested leads in a cross-sectional SNPxAge interaction model are presented in **Supplemental Table 4** and the longitudinal SNPxAge interaction model results are in **Supplemental Table 5**. Summary statistics for the cross-sectional UKB leads tested in MESA are in **Supplemental Table 6 and Supplemental Table 7.**

**Table 3.**
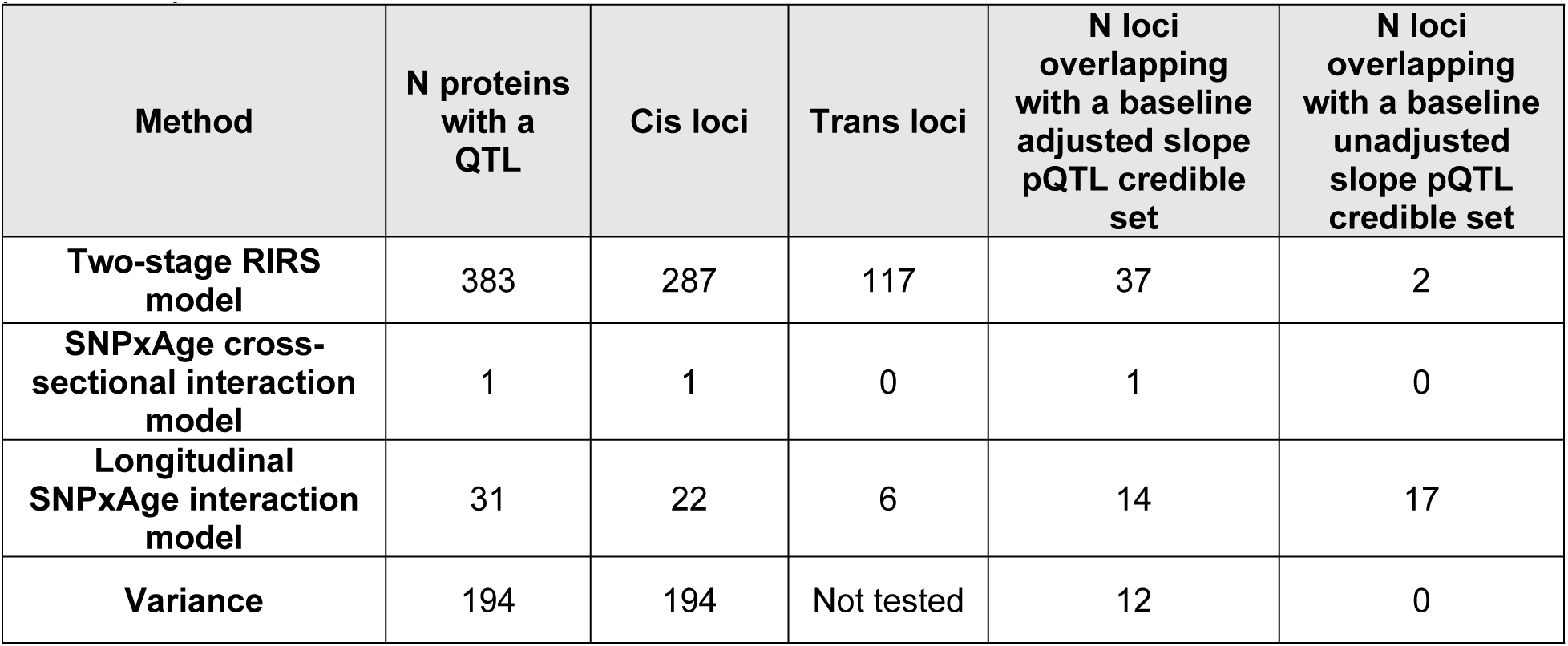
Number of loci identified using different longitudinal and variance modeling methods. For the two stage random intercept random slope (RIRS) model and variance QTL analysis, all 2,737 proteins were tested and a significance threshold of P < 5E-08 was used for cis loci and p < 1.7E-11 was used for trans loci. For the cross-sectional SNPxAge interaction model and longitudinal SNPxAge interaction model, MESA slope leads tested were determined by selecting the variant with the highest posterior inclusion probability (PIP) within each credible set from baseline adjusted and baseline unadjusted analyses; interaction effects were considered significant if P < 6.96E-5. Loci were considered overlapping with a slope pQTL (either baseline adjusted or baseline unadjusted) if any significant variant was also found in a corresponding protein slope credible set.

When using a two-step procedure but with a random slope outcome, 383 proteins had a slope pQTL in MESA (**Table 3, Supplemental Table 8**). Unlike the linear slope baseline adjusted analyses, the majority of the lead variants were directionally discordant with the cross-sectional association, but 30 loci were also found in credible sets in the baseline-adjusted analysis and 9 loci in the baseline unadjusted analysis.

Adjusting for the mean protein level instead of the baseline protein level did not alter slope pQTL discovery however SNP effect directions were negatively correlated with the SNP effect estimates derived from the baseline unadjusted model; and there was no correlation observed between SNP effect sizes derived from models adjusting for mean protein level and models adjusting for the baseline protein level (**Supplemental Figure 3B,C**). Further adjustment for change in eGFR did not impact QTL discovery and beta estimates were similar to the baseline adjusted analysis beta estimates (**Supplemental Figure 4**). There was little evidence of visit heterogeneity with only 23 of the 718 leads tested (0.3%) with a significant Cochran’s Q p-value < 6.96e-5 (**Supplemental Table 9**).

### Overlap with variance QTLs

Of the 2,737 proteins tested in MESA, 194 had a cis variance pQTL. The majority of proteins with a cis vQTL also had either a baseline adjusted or baseline unadjusted slope cis pQTL (147/194, 75.8%). Of those 141, 71 proteins had variance pQTLs which were directionally concordant with the slope pQTL, and 124 were directionally discordant. However, only 15 variance loci contained a variant that that was also included in a corresponding protein slope credible set, (12 from the baseline adjusted analysis and 3 from the baseline unadjusted analysis). Overlap of slope pQTLs with variance QTLs from a better-powered dataset was also investigated.(10) Of the 473 proteins with a cis vQTL identified in Hillary et. al., 258 (59.0%) had a slope pQTL (considering both baseline unadjusted and baseline adjusted analyses) identified in MESA . Sixteen of the lead vQTL variants were also included in a slope pQTL credible set (either baseline unadjusted or baseline adjusted) for the corresponding protein. Variance pQTL summary statistics with lookups from baseline unadjusted and baseline adjusted summary statistics are available in **Supplemental Table 10**.

### Biological characterization of slope loci and proteins

Using FAVOR, the majority of slope leads (considering both baseline unadjusted and baseline adjusted analyses) were in intronic regions (**Supplemental Table 11**). Almost all leads have been previously implicated in various GWAS studies; a complete lookup table is available in **Supplemental Table 12**. Of the 603 proteins with a slope pQTL credible set, 489 were also found to be significantly associated with age in previous work by Sun et. al. in UK Biobank, demonstrating a significant enrichment versus all tested proteins(21) (P = 4.6E-15, Fisher’s Exact Test). We also tested for enrichment of proteins with a slope pQTL with those included in the proteomic aging clock as identified by Argentieri et. al.(3) Of the 240 proteins selected for that clock, 40 have slope pQTLs (p = 0.38, Fisher’s Exact Test).

## Discussion

The dynamic nature of the human proteome has been described,(22,23) but the extent to which these changes are genetically driven is unclear. Understanding the genetic drivers of protein changes over time can have implications in the context of biomarker development and precision medicine. We identified 20 slope pQTLs in MESA in baseline unadjusted analyses and only 4 loci replicated in an external cohort; however our results highlight important modeling considerations and the need for larger sample sizes to identify robust slope loci. To that end, we explored the impact of data transformation on semi-quantitative protein measures and the slope calculation. Second, we conducted a genome-wide scan to identify loci influencing changes in circulating protein levels, with consideration for the impact of adjustment for the baseline protein measure. Third, we compared overlap of slope pQTLs with other measures of protein variability and loci identified using potentially more robust modeling strategies. Finally, our results provide caution on challenges with interpreting covariate adjustment for baseline protein levels in models of slope phenotypes.

### Impact of NPX data transformation on slope

The levels of circulating proteins for all cohorts used in this analysis were measured using the Olink platform, a semi-quantitative assay that returns protein levels in normalized expression (NPX) units; thus a data transformation step is often recommended. Regardless of data transformation, slopes were highly correlated. There were some slope outliers observed when calculating the slope using the NPX values; 94% of proteins had at least one outlier but outliers were still rare—the greatest outlier proportion (with outliers considered to be +/- 5SD from mean slope) was 0.01%. However, with only three samples per individual it is unclear if outliers are reflecting true biology or technical noise. Since the rank inverse normal transformation is frequently used in cross-sectional pQTL analyses(21) and enables inclusion of outliers it was the selected transformation used for the present analyses. It should be noted that the slope calculated from the INT protein values is capturing more so a change in rank than change in protein level, and so a population-wide change in protein level (e.g., everyone has a reduction in circulating IL-3 levels) may not be captured. However, this work aimed to identify genetic drivers of inter-individual differences in protein slopes and thus potentially overlooking these types of associations was not a concern.

### Slope pQTL discovery and influence of baseline adjustment

With a large number of features (nearly 3,000 proteins and millions of SNPs) tested, identifying a method that was computationally feasible, replicable, and did not overfit the data was necessary. Although prior genome-wide studies of focused on a single trait and were thus able to utilize more computationally intensive methods(12,20) those analyses often took several days to run—an approach that quickly becomes computationally infeasible with nearly 3,000 traits. A two-stage approach in which a slope is calculated in the first step and then used as the outcome in a traditional QTL model in the second step enables the analysis to have a similar compute time as a cross-sectional analysis.

The model with baseline adjustment resulted in the greatest number of proteins with a slope pQTL (607 proteins), but when not adjusting for the baseline measure only 19 proteins had a slope pQTL. To date, there is no standard on baseline adjustment for QTL analyses on protein changes; whether to include the baseline measure as a covariate in slope association analyses in general is not an entirely settled matter (although baseline adjustment is frequently employed in a randomized clinical trial setting). However, recent work does strongly encourage the use of baseline unadjusted models.(24,25) Given the strong association between protein baseline value and protein slope, there are two possible paths illustrating the relationship between a SNP, the baseline protein value, and the slope, as illustrated in **Figure 5**. The baseline protein measure could be a mediator, which should often not be adjusted for unless the direct effect only is of interest. In the second case, baseline is a collider. Here, “potential confounders” could be measured technical and environmental effects (such as batch or plate), or even more problematic unmeasured effects. Conditioning on the baseline protein measure would open this path and result in bias unless all potential upstream confounders were conditioned on which would be very difficult, if not impossible. While it is difficult to classify exactly which baseline protein measures are acting as a collider or mediator, we did consider the impact of measured technical effects such as change in eGFR (which does strongly impact circulating protein levels) and batch/plate across visits. Models adjusting for change in eGFR did not attenuate findings. Also, almost all subjects in all three cohorts had all proteomics samples assayed on the same plate, reducing measurement error attributed to those technical effects. It is also possible that given the expected correlation between the slope and baseline measure, spurious associations between the SNP and protein slope could arise due to regression to the mean.(26) In the baseline adjusted analysis, the betas were consistently larger and standard errors consistently smaller compared to the baseline unadjusted analysis. It may also be possible that many of the proteins measured do not change over the time period captured or may be more impacted by time-limited exposures. Protein measures over shorter time periods (2-5 years) have been shown to be relatively stable but some proteins to exhibit significant variability.(7,23,27) However, whether these phenomenon are biasing results to the null or inducing spurious associations is unclear.

With the potential bias introduced by baseline adjustment, we caution adjustment for the baseline measure in genetic analyses of protein change, but as recommended by Glymour et. al. and implemented in Gorski et. al., we present both baseline adjusted and unadjusted results and consider slope pQTLs identified in both models. Unlike the work of Gorski et. al. which focused on eGFR decline, the nearly 3,000 proteins tested do not all have a “ground truth” rate of change which makes determining whether the baseline protein measure could be a collider difficult. Although the baseline adjusted analysis identified several hundred more slope loci, we hypothesized that the likely true associations would be directionally consistent with effect sizes from baseline unadjusted analyses and thus also have a higher replication rate; and associations that were directionally discordant between baseline adjusted and baseline unadjusted analyses would likely reflect regression to the mean effects and have lower replication rates. In the baseline adjusted analyses, the majority of loci were only significant when adjusting for the baseline measure—only five credible sets contained a variant that was also significant in baseline unadjusted analyses, and in two of those credible sets the estimate of the corresponding SNP in baseline unadjusted analyses was in the opposite effect direction. The majority of credible sets were significant in the baseline adjusted analyses only, and directionally discordant with the baseline unadjusted estimate. This category of credible sets also had the highest replication rate in CARDIA and UKB. Contrary to our hypothesis, credible sets significant in baseline adjusted analyses only and directionally concordant with baseline unadjusted analyses had much lower replication rates. This demonstrates that the bias introduced by adjusting for the baseline measure is not cohort-specific and signals should be interpreted with caution, regardless of successful replication and directional concordance with cross-sectional estimates. Summary statistics for both baseline adjusted and baseline unadjusted analyses are provided and researchers should consider which model is most appropriate for their use; noting that prior studies generally suggest baseline unadjusted models.(24,26,28)

### Comparison to other modeling strategies of protein change and variance

Given the method used to identify slope pQTLs is an attempt to approximate a longitudinal model to reduce computational burden, we investigated if slope pQTLs were discoverable using other more robust methods. A two-step random intercept random slope model has been found to have adequate Type I Error and power (but large bias) in prior work.(14) However, we found that the SNP effect sizes obtained with the random slope outcome were largely discordant with the cross-sectional effect, suggesting that this method also yields associations potentially due to regression to the mean effects. Although a genome-wide scan using a two-visit slope was not conducted, given the high correlation with three-visit slopes (Pearson correlation coefficient = 0.99) we anticipate the same loci would be identified.

Prior researchers have hypothesized that SNPxAge interactions may overlap with slope loci.(29) However, we found almost no overlap with slope pQTL leads (or UKB cross-sectional leads, see **Supplemental Table 6**) and cross-sectional SNPxAge interaction effects. When utilizing the longitudinal data in a longitudinal SNPxAge interaction model, only 31 slope pQTL leads (of the 630 leads tested) had a significant interaction effect; however 89% of the baseline unadjusted leads (vs. 0.02% of baseline adjusted leads) did have a significant interaction effect suggesting that not adjusting for the baseline may better proxy true longitudinal effects. However, the majority of interaction effects were directionally discordant with the corresponding main effect suggesting regression to the mean may be present. Both cross-sectional and longitudinal interaction analyses utilized the same subset of participants as in the slope pQTL analysis; thus we may be underpowered to detect interaction effects. Future studies in larger cohorts, advances in longitudinal modeling, and improvements in computational hardware may enable increased power and efficiency to detect slope effects using linear mixed modeling approaches.

It is possible that protein variability (driven by either genetic or non-genetic factors) could influence discovery of slope pQTLs. To assess this, we investigated the overlap of protein variance QTLs with slope pQTLs. Many of the proteins with a variance QTL also had a slope QTLs (but with the baseline adjusted model only). There was no apparent pattern in whether proteins with slope pQTLs tended to also have variance increasing or decreasing QTLs as effect directionality was relatively evenly split. This trend is also present when comparing overlap with vQTLs identified in UK Biobank, where 230 cis vQTLs with a slope pQTL are variance increasing, and 203 are variance decreasing. Although slope pQTL signals largely did not overlap with variance signals, there was an enrichment for proteins with a variance QTL also having a slope pQTL. Part of this enrichment may be reflecting the possibility that many of the baseline adjusted slope QTLs may be due to collider bias (as discussed above) rather than true slope associations; and there is a large overlap of main effect pQTLs and variance pQTLs (over 90% overlap identified in UKB(10)). For example, positive slope loci that are also variance increasing loci might indicate that the slope is capturing measurement error and/or an environmental interaction, leading to a spurious association. Slope loci with discordant effect directions with the variance effect (i.e., the locus is associated with decreasing variance but increasing slope) could indicate a true slope effect. Of proteins with both a variance QTL and slope QTL, the majority had a positive slope effect but negative variance effect.

### Potential biological implications

Many of the loci identified may have biological implications. Of the annotated credible set leads, approximately 18% are in exonic protein coding regions. Of the 718 total slope credible sets, many are at relevant loci for complex traits. For example, when looking up leads in the GWAS Catalog,(30) variants in credible sets for the slope of SSC5D, LRPAP1, HCG22, SHBG, and APOE have been previously associated with waist-to-hip ratio adjusted for BMI, and variants in credible sets for 40 proteins have been implicated in various measures of cholesterol levels. We found significant enrichment for age-associated proteins with a baseline adjusted slope pQTL, providing some support that the slope pQTLs are capturing proteins known to change over time. However, we found minimal overlap of proteins with slope QTLs and proteins included in a recent proteomic aging clock (also created in UK Biobank). Selection of proteins for aging clocks can be sensitive to the highly correlated nature of proteomics data, so the minimal overlap is not entirely unexpected and we caution overinterpretation.

### Limitations

There exists a relatively limited literature on slope QTLs, especially in the context of molecular traits such as proteins. The present work provides a preliminary scan of a type of molecular slope QTLs, compares multiple modeling strategies including the impact of protein data normalization, slope calculation, and model specification. However, this work also has many limitations. As noted previously, the protein values were rank inverse normal transformed prior to the slope calculation; as such the resulting slopes were capturing a change in rank within the population rather than a true change in circulating protein level. Thus, it is possible we were unable to detect genetic loci associated with population-wide changes in protein levels. The slope measure used assumes linearity, but this assumption may not be valid for all proteins. Future work should explore non-linear models and analytical approaches, especially when larger sample sizes and more timepoints are available. Collapsing the three measures into a slope summary statistic also means that some of the value of having multiple measures per individual is lost, although with the relatively small sample size in MESA we had limited power to detect effects using a linear mixed model. Additionally, the inconsistent time gaps between proteomics timepoints means that the change over some periods has a greater impact on the slope. Sample and cohort differences could also impact slope pQTL discovery and replication. Although the proteomics visits in all three cohorts spanned middle age, the average age at baseline within each cohort varied. Different physiological changes may be taking place, which could impact ability to detect protein slope QTLs. All cohorts did also use different versions of the Olink assay which could impact replicability.(31) Additionally, the analysis sample is also likely to be healthier than average (since we select for individuals who survived and attended later study visits). Finally, many environmental changes impacting circulating protein levels likely took place during the time span covered in each cohort (such as medication use and disease), and it is not possible to account for all of them.

The present analysis represents one of the first attempts to identify genetic drivers of protein changes over time. Our results highlight considerations for modeling strategies of change phenotypes and build on understanding of potential genetic mechanisms influencing interindividual proteome changes over time.

## Methods

### Cohorts

The Multi-Ethnic Study for Atherosclerosis (MESA) study is a population-based cohort study of 6,814 participants aged 45 to 84 years old recruited from six sites across the US(32). All participants were free of clinical CVD at baseline. MESA is a diverse cohort with 38% of participants self-identifying as White, 28% as African American, 22% as Hispanic, and 12% as Asian. After removing related participants, 2,007 participants had longitudinal proteomics data spanning ∼18 years from Exam 1 (2000-2002), Exam 5 (2010-2012) and Exam 6 (2016-2018) and were used for analysis. The Coronary Artery Risk Development in Young Adults (CARDIA) study is a population-based cohort study of 5,115 Black and White participants aged 18-30 years old recruited from four sites (Birmingham AL, Chicago IL, Minneapolis MN, and Oakland CA) starting in 1985-1986(33,34). After removing related participants, 1328 participants have longitudinal proteomics data spanning ∼15 years from Year 15 (2000-2001), Year 20 (2005-2006), Year 25 (2010-2011), and Year 30 (2015-2016). All individuals in the TOPMed cohorts have WGS generated from TOPMed Freeze 10 data(35) (average sequencing depth > 30X), with methods previously described. Pass variants were filtered to MAF > 0.05. For both cohorts, related individuals (n = 112 removed for MESA and n = 5 removed for CARDIA) were identified using KING and relatives of second-degree relatedness were removed using plink. All MESA and CARDIA participants included in this analysis provided written informed consent for use of genetic and multi-omics data, and both studies were approved by the Institutional Review Boards of participating institutions. The authors did not have access to information that could identify individual participants during or after data collection, and data were accessed January 2025.

The UK Biobank (UKB) is a population-based cohort of about 500,000 participants from the United Kingdom(36). Approximately 1,200 participants have longitudinal proteomics data at three visits, with the baseline exam occurring in 2006-2010, and follow-up exams in 2014 and 2019(21). The proteomics data released to the research community has undergone various QC steps as described in Sun et. al. Only participants of European ancestry (as identified using k-means clustering as in(37)) were included for a sample size of 947. Genotyping and imputation has been described previously.(36) Imputed genetic variants were filtered for MAF > 0.05 and INFO > 0.7. Variants were mapped to the hg38 build using the mapping files available in Synapse from Sun et. al.(21) Data were accessed October 2023, and all UK Biobank participants included in analysis provided written informed consent.

### Proteomic data processing

Olink proteomics data was obtained from EDTA plasma samples from the two TOPMed studies (MESA and CARDIA) during at least three separate visits(38,39). Briefly, the Olink platforms used (Olink Explore 3k and Olink Explore HT for MESA and CARDIA respectively) capture approximately 3,000 – 5,000 proteins via a proximity-extension assay which uses antibody-oligo complexes to recognize and bind target proteins(40). The resulting relative protein abundances are given in normalized protein expression (NPX) units. In general, samples from the same subject in both cohorts were assayed in the same batch/plate, reducing technical noise that could arise due to samples from different visits (but the same subject) being assayed in different batches/plates. In MESA, 0.03% of subjects had at least one sample assayed on a different plate; in CARDIA no subjects had a sample assayed on a different plate. In UKB, baseline proteomics measures were assayed using the Olink 3k platform while follow-up exams were assayed on the Olink 1.5k platform, enabling longitudinal assessment of 1,474 proteins(41). As a result, all UKB subjects have at least one sample assayed in a different batch or plate than the others. Within each cohort (MESA, CARDIA, and UKB) proteins with high rates of missingness (>50%) within and across batches and intra-plate coefficients of variation > 20% were dropped from analysis (**Figure 6, Supplementary Table 1**). Protein principal components were estimated; samples that were 5 standard deviations from the mean of protein PC1 and PC2 were also excluded.

**Figure 6.**
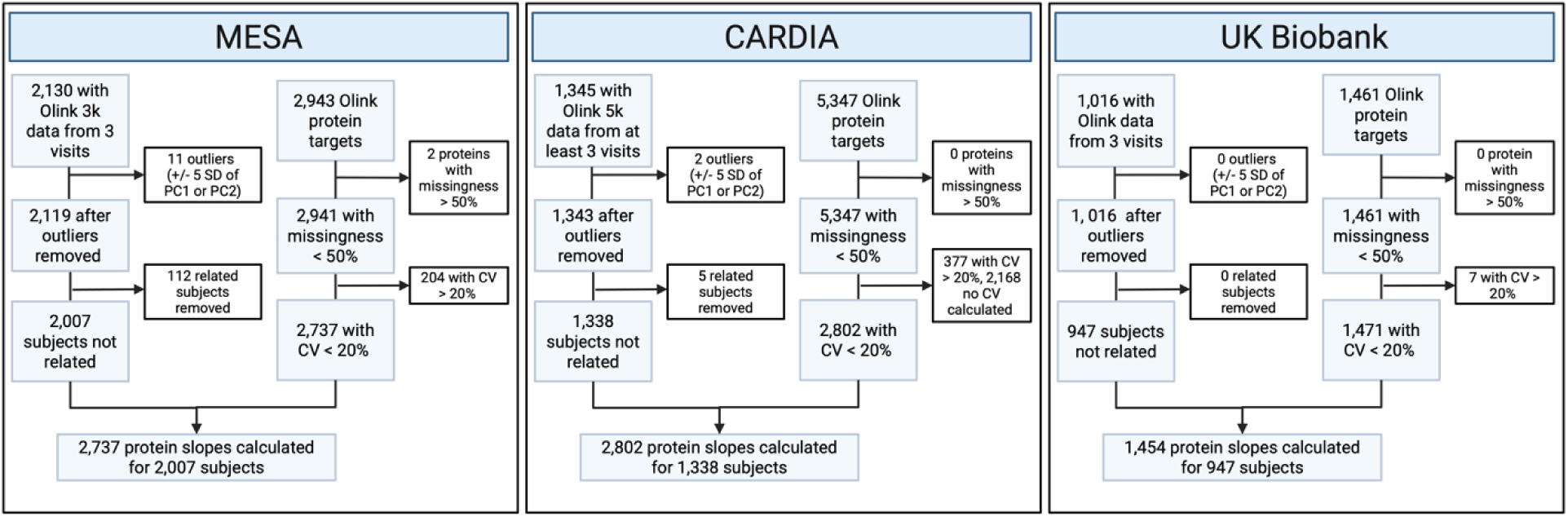
Flowchart illustrating cohort and protein selection. Proteins were considered to have a high rate of missingness if the missingness percentage exceeded 50% within and across batches (if batches were used); in all cohorts missingness rates were very low. Samples were considered outliers if they fell outside five standard deviations from the mean of PC1 or PC2. Subjects were included in analysis if they had a proteomics sample that passed the prior QC steps from at least three separate timepoints. A subset of unrelated subjects was identified using a kinship coefficient of 0.0884 (second-degree relatedness). Figure created using Biorender.

Given the semi-quantitative nature of Olink measures, generally data transformation is recommended. The slope calculation may be sensitive to the data transformation used; as such three commonly used data transformations were applied prior to the slope calculation and the resulting slopes were compared using MESA data only. First, protein measures were standardized (to mean=0 and standard deviation=1) and rank inverse normal transformed within each cohort and visit (similar to prior work).(21,39) Second, protein residuals were calculated by regressing the protein values on age, age squared, sex, site, and ten ancestry PCs. Finally, no transformation was applied and a slope was calculated using the NPX values. Pearson’s correlations between the resulting slopes calculated from each transformation were compared.

### Slope calculation: linear model

Within each cohort and protein, a linear model for each participant with the protein measures as the dependent variable and age as the independent variable estimated. The resulting age coefficient (representing the 1-SD change per year of the protein for a given participant) was extracted. These per-person per-protein slopes were then rank-inverse normal transformed. Since the proteomics data was not imputed, some participants have missing measures for some proteins; thus the total N analyzed per protein varied slightly. Pearson correlation coefficients between slope values and cross-sectional values were calculating to understand any correlation between the measures.

### QTL analysis and fine-mapping

The resulting protein slopes were used as the outcome in QTL analyses. All QTL analyses were run using TensorQTL(42). Since TensorQTL does not allow for inclusion of a kinship matrix, individuals with second-degree relatedness were identified using KING and analysis samples containing unrelated individuals were selected (kinship coefficient = 0.0884). For the linear slope, QTL analyses were adjusted for baseline age, baseline age squared, sex, the first ten ancestry principal components (PCs). Analyses with further adjustment for the baseline measurement were also completed. When necessary, additional study-specific covariates were included: study site/center (MESA and CARDIA) and genotype chip (UKB only). Proteomics batch or plate were not adjusted for in the MESA and CARDIA analyses as all samples for each subject were assayed within the same batch/plate. Slope cis-pQTLs were defined as variants with P < 5E-08 within 1 megabase (Mb) of the transcription start side (TSS) and slope trans-pQTLs were defined as variants further than 1Mb from the TSS and P < 1.83E-11 [5E-08/2737 proteins tested]. SuSiE(43) was used to fine-map significant loci using individual-level LD. Credible sets without any genome-wide significant SNPs (P < 5E-08) were filtered out.

For both the baseline adjusted and baseline unadjusted analyses in MESA, slope QTLs were grouped into four categories based on replication in either the baseline adjusted or baseline unadjusted model and effect direction concordance. A baseline unadjusted QTL was considered overlapping if any variant in the credible set had P < 5E-08 in the baseline adjusted analysis (and vice versa). To determine replication in external cohorts, variants from MESA slope credible sets were looked up in CARDIA and UKB. If any variant within a credible set had P < 5.31E-06 (Bonferroni-adjusted threshold based on the total number of MESA baseline unadjusted and baseline adjusted credible set SNPs) and effect direction concordance in the external cohort, the slope QTL was considered replicated. Not all variants tested in MESA were tested in CARDIA and UKB due to population allele frequency differences (all analyses filtered for MAF > 0.05).

### Cross-sectional QTL analysis

To assess the overlap of cross-sectional and slope associations within the same sample, the MESA baseline (Visit 1) protein measures from the same n = 2,007 subset were used in a cross-sectional pQTL analysis. Models adjusted for age, sex, site, and 10 ancestry PCs. Similar to the slope pQTL analysis, cis-pQTLs were defined as variants with p < 5E-08 within 1 megabase (Mb) of the transcription start side (TSS) and trans-pQTLs were defined as variants further than 1Mb from the TSS and p < 1.83E-11 [5E-08/2737 proteins tested]. Slope credible set variants were considered overlapping with cross-sectional signals if the variant had p < 5E-08 in the cross-sectional analysis.

### Sensitivity analyses and comparison to other longitudinal methods: Two-stage random intercept random slope model, SNPxAge interaction model, longitudinal SNPxAge interaction model, and variance pQTLs

Four additional modeling strategies were implemented using MESA to explore if any loci identified using these methods overlapped with either baseline adjusted or baseline unadjusted slope pQTLs. Unless specified otherwise, only lead variants from both baseline unadjusted and baseline adjusted analyses were tested. Leads were defined as the variant with the highest posterior inclusion probability (PIP) within each credible set; resulting in a total of 718 leads tested.

Prior work has hypothesized that genetic variants influencing change of a trait are a subset of genetic variants with a cross-sectional SNPxAge interaction.(29). Lead SNPs were tested in a cross-sectional SNPxAge interaction model using the MESA visit 1 measures. Models adjusted for age, sex, site, and ten ancestry PCs. Interaction effects were considered significant based on adjustment for the SNP-protein pairs tested (P < 6.96-E05 [0.05/718]).

To determine if a longitudinal SNPxAge model would reveal the same associations, we tested lead SNPs in a longitudinal SNPxAge interaction model with the R package lmerTest(44). The model included fixed terms for the SNP, sex, site and ten ancestry PCs; a random intercept and random slope for age; and an interaction term for SNP and age. Ages were centered at 50 and divided by 10 for appropriate scaling and model convergence.(14) Interaction effects were considered significant based on adjustment for the SNP-protein pairs tested (P < 6.96E-05 [0.05/718]). Leads as identified in the Sun et. al. cross-sectional UKB pQTL analysis were also tested; for further details see **Supplemental Text**.

Other studies of longitudinal trait change have utilized a two-stage random intercept random slope model for computational efficiency.(14) In the first step, a linear mixed effects model adjusting for age (centered at 50 and divided by 10 for appropriate scaling and model convergence(14)), sex, site, and a random intercept and random slope for participant was calculated for each protein (the dependent variable). The per-person random slopes were extracted and in step two used as the outcome in a genome-wide pQTL scan with adjustment for ancestry PCs. Lead variants were determined using distance-based (1Mb) clumping. Leads as identified in the Sun et. al. cross-sectional UKB pQTL analysis were also tested; for further details see **Supplemental Text**.

Variance pQTLs were identified following a procedure outlined in previous work with UK Biobank.(10) First, protein residuals from the baseline visit were calculated by regressing the protein value on age, age squared, sex, site, and ten ancestry PCs. Levene’s test (using median) was performed on *cis* regions for the 2,737 proteins using OSCA which calculates effect sizes from the z-statistic.(45,46) The same n = 2,007 sample used for the slope analysis was also used for the variance QTL analysis. Significant loci (p < 5E-08) were identified using distance-based clumping. Genome-wide scans for each protein were not tested due to computational burden; only cis loci were tested and identified. Variants within slope pQTL credible sets (either baseline adjusted or baseline unadjusted) with a p < 5E-08 in the variance QTL analysis were considered overlapping. Lookups of baseline-adjusted and baseline unadjusted slope pQTL credible sets in UKB variance QTLs were also conducted (also using a threshold of p < 5E-08).

Additional sensitivity analyses conducted in MESA further adjusted for either the mean protein level across three visits or the change in eGFR between visit 1 and visit 6. For both visits eGFR was calculated using the 2021 CKD-EPI equation using serum creatinine measured at each visit. For the subset of both baseline adjusted and baseline unadjusted protein slope credible set leads, heterogeneity of the SNP effect on protein levels across visits was also assessed using Cochran’s Q and I^2^ statistics.

### Variant effects and GWAS associations

The predicted effects of lead credible set variants (defined as the variant with the highest posterior inclusion probability (PIP) within each credible set of both baseline adjusted and baseline unadjusted analyses) were determined using FAVOR.(47) Overlap with GWAS variants was determined by looking up lead credible set variants in the GWAS Catalog(30) (accessed April 2025). Proteins with slope pQTLs (baseline adjusted and baseline unadjusted) were also compared to age-associated proteins as identified in Sun et. al.(21)

## Acknowledgements

Molecular data for the Trans-Omics in Precision Medicine (TOPMed) program was supported by the National Heart, Lung and Blood Institute (NHLBI). Whole Genome Sequencing (WGS) for “NHLBI TOPMed: Multi-Ethnic Study of Atherosclerosis (MESA)” (phs001416.v1.p1) was performed at the Broad Institute of MIT and Harvard (3U54HG003067-13S1). WGS for “NHLBI TOPMed: Coronary Artery Risk Development in Young Adults (CARDIA)” (phs001612.v3.p3) was performed at Baylor College of Medicine Human Genome Sequencing Center (HHSN268201600033I). Core support including centralized genomic read mapping and genotype calling, along with variant quality metrics and filtering were provided by the TOPMed Informatics Research Center (3R01HL-117626-02S1; contract HHSN268201800002I). Core support including phenotype harmonization, data management, sample-identity QC, and general program coordination were provided by the TOPMed Data Coordinating Center (R01HL-120393; U01HL-120393; contract HHSN268201800001I). We gratefully acknowledge the studies and participants who provided biological samples and data for TOPMed.

MESA and the MESA SHARe project are conducted and supported by the National Heart, Lung, and Blood Institute (NHLBI) in collaboration with MESA investigators. Support for MESA is provided by contracts 75N92025D00022, 75N92020D00001, HHSN268201500003I, N01-HC-95159, 75N92025D00026, 75N92020D00005, N01-HC-95160, 75N92020D00002, N01-HC-95161, 75N92025D00024, 75N92020D00003, N01-HC-95162, 75N92025D00027, 75N92020D00006, N01-HC-95163, 75N92025D00025, 75N92020D00004, N01-HC-95164, 75N92025D00028, 75N92020D00007, N01-HC-95165, N01-HC-95166, N01-HC-95167, N01-HC-95168, N01-HC-95169, UL1-TR-000040, UL1-TR-001079, UL1-TR-001420, UL1TR001881, and R01HL105756. The authors thank the MESA participants and the MESA investigators and staff for their valuable contributions. A full list of participating MESA investigators and institutions can be found at http://www.mesa-nhlbi.org.

The Coronary Artery Risk Development in Young Adults Study (CARDIA) is conducted and supported by the National Heart, Lung, and Blood Institute (NHLBI) in collaboration with the University of Alabama at Birmingham (HHSN268201800005I & HHSN268201800007I), Northwestern University (HHSN268201800003I), University of Minnesota (HHSN268201800006I), and Kaiser Foundation Research Institute (HHSN268201800004I). CARDIA was also partially supported by the Intramural Research Program of the National Institute on Aging (NIA) and an intra-agency agreement between NIA and NHLBI (AG0005).

This research has been conducted using the UKB Resource under Application Number 25953. We thank the UKB participants and research team for enabling this study.

